# Walking cadence affects the recruitment of the medial-lateral balance mechanisms

**DOI:** 10.1101/658070

**Authors:** Tyler Fettrow, Hendrik Reimann, David Grenet, Jeremy Crenshaw, Jill Higginson, John Jeka

## Abstract

We have previously identified three balance mechanisms that young healthy adults use to maintain balance while walking. The three mechanisms are: 1) The lateral ankle mechanism, an active modulation of ankle inversion/eversion in stance; 2) The foot placement mechanism, an active shift of the swing foot placement; and 3) The push-off mechanism, an active modulation of the ankle plantarflexion angle during double stance. Here we seek to determine whether there are changes in neural control of balance when walking at different cadences and speeds. Twenty-one healthy young adults walked on a self-paced treadmill while immersed in a 3D virtual reality cave, and periodically received balance perturbations (bipolar galvanic vestibular stimulation) eliciting a perceived fall to the side. Subjects were instructed to match two cadences specified by a metronome, 110bpm (*High*) and 80bpm (*Low*), which led to faster and slower gait speeds, respectively. The results indicate that subjects altered the use of the balance mechanisms at different cadences. The lateral ankle mechanism was used more in the *Low* condition, while the foot placement mechanism was used more in the *High* condition. There was no difference in the use of the push-off mechanism between cadence conditions. These results suggest that neural control of balance is altered when gait characteristics such as cadence change, suggesting a flexible balance response that is sensitive to the constraints of the gait cycle. We speculate that the use of the balance mechanisms may be a factor resulting in well-known characteristics of gait in populations with compromised balance control, such as slower gait speed in older adults or higher cadence in people with Parkinson’s disease.

## 1. Introduction

There is an ongoing debate about whether or not walking slower is more stable (Bruijn et al., 2009). We know certain patient populations reduce their gait speed and increase their cadence (Buckley et al., 2018; Duan-Porter et al., 2019; Himann et al., 1988; Lauretani et al., 2003), but is the motivation to improve stability? Key factors that lead to the decrease in gait speed in older adults remains unresolved, but we speculate that the use of the control of balance plays a larger role than previously recognized. Our focus here is to assess changes in the control of balance at different gait speeds, and more specifically cadence.

There is support for slower walking being more stable. Though a firm definition is not established, stability has been quantified with different methods. Roos and Dingwell (2013) found that neuromuscular noise is diminished when walking slower. Less noise generally results in better estimates of a physical quantity, and thus improved control. Lyapunov exponents, a common tool used for quantifying overall stability, have been shown to be reduced at lower walking speeds (Dingwell and Marin, 2006; England and Granata, 2007). Others have pointed to neurological deficiencies associated with slower walking, without referring to stability directly. Anson et al. (2018) have shown vestibular function loss causes people to walk with longer, slower steps, and Menz et al. (2004) have shown people with peripheral neuropathy reduced walking speed and cadence. Hsiao et al. (2017) suggested that a lateral weight shift mechanism is impaired in chronic stroke, leading to reduced gait speed.

Another body of literature suggests fast walking is more stable. Variability, which is often attributed to greater instability, lessens with increased velocity (Donker and Beek, 2002). Studies have also used the mediolateral margin of stability to quantify overall stability, which increases with speed (Hof et al., 2007; Gates et al., 2013). A larger margin of stability is viewed as safer and more stable. Recent modeling results indicate that slowed gait can be explained entirely by diminished muscle strength (Song and Geyer, 2018). However, Fan et al. (2016) has shown older adults with slower gait can walk faster if instructed, suggesting that locomotor capacity (i.e. force-generating capabilities) are not the reason for slowed gait. This suggests a complex relationship between stability and slowed gait, as muscle strength is considered necessary but its role in the control of balance is unknown, with other factors such as multisensory integration playing an important role (Peterka, 2002; Oie et al., 2002).

Regardless of the walking speed, a common theme to the process of keeping a body upright is the relationship between the center of mass (CoM) and the center of pressure (CoP). The behavior of the CoM behavior, can be explained by the relationship between CoP and CoM, as in Equation 1 (Winter, 1995).

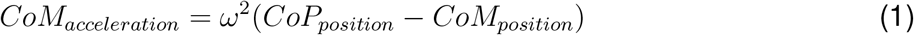

The CoM acceleration is proportional to the distance between the CoP and CoM. Where *ω* = *g*/*l*, l is the height of the CoM above ground and g the acceleration from gravity. This relationship holds true for both the anterior-posterior direction and the medial-lateral direction.

Applying Equation 1, a threat to balance would be an undesired movement of the CoM relative to the CoP. To correct this undesired movement, a motor action has to be taken to accelerate the CoM in the opposite direction or, rather, to decelerate the CoM, thus stopping the movement. To prevent a fall, the CoM should not be able to accelerate to the point of no return in any particular direction. So the CoP must be moved to accelerate the CoM in the opposite direction of its current travel.

In a previous study with young healthy adults, we have identified three different mechanisms of balance control in response to perceived falls (Reimann et al., 2018b). The perceived fall theoretically induces a perceived shift of the center of mass (CoM), requiring a motor action that shifts the CoM in the opposite direction. The following mechanisms are distinct balance mechanisms, observed in previous experiments, that can shift the CoP and CoM in the medial-lateral direction.

The foot placement is the most commonly reported and quantified method of balance control (Kuo, 1999; Bauby and Kuo, 2000; Vlutters et al., 2016). The foot placement mechanism consists of the swing foot moving in the direction of the perceived fall, and on heel strike, shifts the net CoP in the direction of the perceived fall. In general, the foot placement mechanism refers to the modulation of step width in a particular direction (i.e. the direction of the perceived fall). Wang and Srinivasan (2014) show that 80% of the foot placement can be explained by the position and velocity of the CoM. Theoretically, the sensory input we provide to subjects alters the perceived state of the CoM, thus there should be a difference between the predicted foot placement based on the CoM behavior, and the actual foot placement. We refer to this measure as the model-corrected foot placement. The lateral ankle is another mechanism that can shift the CoP (Hof et al., 2010). The lateral ankle mechanism refers to the generation of ankle inversion/eversion torque during the single stance phase. Activating musculature to roll the ankle while the foot is on the ground shifts the CoP under the foot in the direction of the perceived fall. The goal is to accelerate the CoM away from the direction of the perceived fall, and by shifting the CoP in the direction of the perceived fall, the CoM will accelerate away from the perceived fall. We refer to the third balance mechanism as the push-off mechanism. The push-off mechanism refers to the modulation of ankle plantarflexion angle during double stance. In response to a visually perceived fall to the side, we observed a direction-dependent modulation of the stance leg ankle plantar/dorsiflexion angle. Very few studies have recognized the push-off mechanism’s role in control of balance in the medial-lateral direction. Ankle plantarflexion torque has been shown to change as a result of bipolar, binaural galvanic vestibular stimulation (Iles et al., 2007). Only recently has the push-off been verified to have a functional role in balance control in the medial-lateral direction as Kim and Collins (2015) show modulation of ankle torque based on CoM behavior can reduce metabolic expenditure. Furthermore, Klemetti et al. (2014) have provided evidence that ankle plantarflexion torque induces trunk roll accelerations, providing further support for the push-off acting in the medial-lateral direction.

The need for multiple mechanisms to promote balance derives from the gait cycle, which demands different mechanisms due to the changing configuration of the body (i.e., alternation of double stance and single stance). Importantly, these mechanisms have a temporal order. The earliest response to a balance disturbance (i.e., at heel strike) is the lateral ankle mechanism, followed by a change in foot placement and push-off (Reimann et al., 2018b). Given that temporal order, we were interested in determining if gait characteristics can change how these balance mechanisms are used. Specifically, are there changes in the use of the balance mechanisms at different cadences? Modeling results suggest more frequent steps leads to more opportunities to correct undesired CoM movements with the foot placement mechanism (Reimann et al., 2018a). We hypothesized that due to the longer single stance in slower walking, the lateral ankle mechanism would play a larger role in balance, and in faster walking, the foot placement mechanism would provide the majority of balance control.

## 2. Methods

### 2.1. Subject Characteristics

Twenty-one healthy young subjects (15 female, 23.65 ± 4.43 years, 1.68 ± 0.11 m, 64.23 ± 14.97 kg) volunteered for the study. Subjects provided informed verbal and written consent to participate. Subjects did not have a history of neurological disorders or surgical procedures involving legs, spine or head. The experiment was approved by the University of Delaware Institutional Review Board.

### 2.2. Experimental design

Subjects walked on a split-belt, instrumented treadmill within a virtual environment projected onto a curved screen surrounding the treadmill as shown in Figure 1 (Bertec Inc., Columbus, Ohio, USA). The treadmill was self-paced, using a nonlinear PD-controller in Labview (National instruments Inc., Austin, TX, USA) to keep the middle of the posterior superior iliac spine (MP-SIS) markers on the mid-line of the treadmill. The same speed command was sent to each belt of the treadmill. The virtual environment consisted of a tiled marble floor with floating cubes randomly distributed in a volume 0-10 m above the floor, 2-17 m to each side from the midline, and infinitely into the distance, forming a 4 m wide corridor for the subjects to walk through (see Figure 1), implemented in Unity3d (Unity Technologies, San Francisco, CA, USA). Perspective in the virtual world was linked to the midpoint between the two markers on the subject’s temples, superposed over forward motion defined by the treadmill speed.

**Figure 1:**
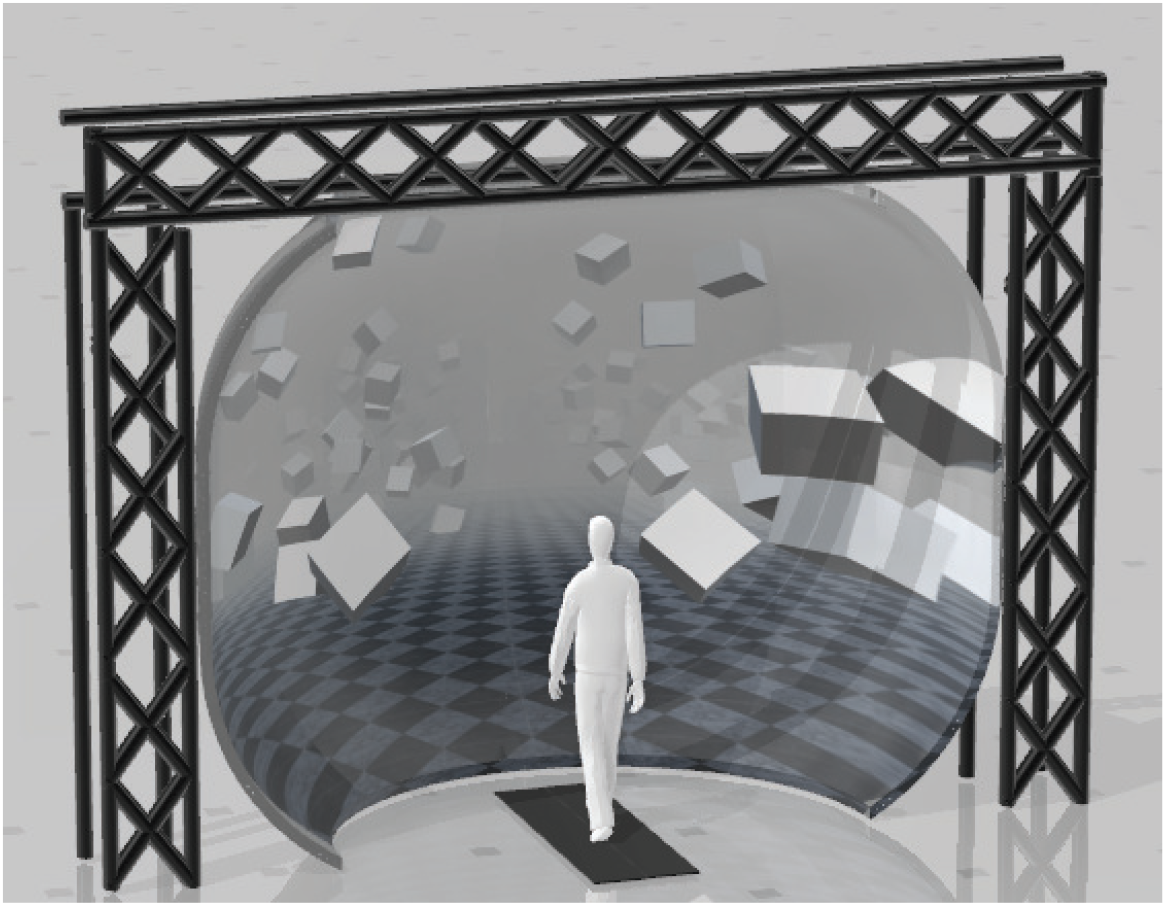
Experimental setup displaying the position of the split-belt treadmill in front of the curved screen.

The subjects performed 20 two-minute trials. A metronome generated by custom Lab-view software played throughout the trials at a frequency of 80 bpm (*Low*) or 110 bpm (*High*), which alternated on every two minute trial. Subjects were asked to match their cadence to the metronome. After a two-minute trial there was a 15 second washout period of no metronome, followed by a 30 second adaptation period for the new metronome frequency before introducing the balance perturbations. Breaks were offered after every 5 two-minute trials.

During the two-minute trials, a target step number was randomized between 10-13. The custom Labview software counted heel strikes until the step counter matched the target step number. The leg of the heel strike that occurs on the target step number is referred to as the trigger leg. The Labview software randomized between two conditions on a trigger, either perturbation or control. Subjects were unaware if the control condition was triggered. The perturbations consisted of a 1 mA current produced by a NeuroConn DC-Stimulator Plus direct current stimulator (Ilmenau, Germany), passed between Axelgaard PALS 3.2cm round electrodes (California, USA) placed on the mastoid processes for 1000 ms, inducing a feeling of falling to the side. The polarity was chosen so that the perturbation always produced the perception of falling towards the trigger leg (i.e. Left heel strikes triggers perturbation, stimulus induced perception of falling to the left).

We identified heel strikes on-line with a manually set vertical threshold of the heel position. The vertical threshold was set to the heel marker position during quiet standing. However, in the *Low* condition people tended to “drag” their feet, which would cause steps to be counted both midway through swing and actual heel strike. To avoid double counting of steps, the experimenter increased the vertical threshold until double counting of steps did not occur.

### 2.3. Data analysis

We recorded full-body kinematics using the Plug-in Gait marker set (Davis et al., 1991), with six additional markers on the anterior thigh, anterior tibia, and 5th metatarsal of each foot. Another additional six markers were placed on the medial femoral epicondyles, medial malleolus, and the tip of the first distal phalanx of the foot for the static calibration pose at the beginning of the collection. We recorded electromyography signals from the tibialis anterior, peroneous longus, medial gastrocnemius, rectus femoris, biceps femoris, gluteus medius, tensor fascia latae, and erector spinae, bilaterally with Cometa’s electromyography PicoEMG sensors (Bareggio, Italy) with hydrogel 30mm x 24mm Covidien Kendall electrodes using SENIAM guidelines for placement (Hermens et al., 2000). Marker positions were recorded at 200 Hz using a Qualisys Miqus motion capture system (Gothenburg, Sweden) with 13 cameras. Electromyography was recorded at 2000 Hz. Ground reaction forces and moments were collected at 1000 Hz.

We used custom Matlab scripts for data management and organization. Force plate data was low pass filtered with a 4th order Butterworth zero-phase filter at a cut-off frequency of 20Hz. EMG data was bandpass filtered with a 4th order Butterworth filter with cutoff frequencies of 20Hz and 500Hz, then rectified, then low-pass filtered with a 4th order Butterworth filter at a cut-off frequency of 6Hz. For each subject and EMG channel, we calculated the average activation across all control strides and used this value to normalize EMG before averaging across subjects. From the marker data, we calculated joint angle data and center of mass (CoM) positions based on a geometric model with 15 segments (pelvis, torso, head, thighs, lower legs, feet, upper arms, forearms, hands) and 38 degrees of freedom (DoF) in OpenSim (John et al., 2007; Seth et al., 2018) using an existing base model (Zajac et al., 1990).

The experimental design allowed for four distinct stimulus conditions, where each leg could trigger a stimulus towards that leg during the 80bpm metronome trials (*Low* cadence) or 100bpm metronome trials (*High* cadence). Here we combine the conditions that are anatomically similar (i.e. [Right heel strike triggers Stimulus Right, Left heel strike triggers Stimulus Left]). This leaves two conditions to analyze, stimulus towards trigger leg in *Low* and *High* cadence conditions. After post-processing, we were left with 1178 perturbations for *Low* cadence and 1567 perturbations for *High* cadence conditions.

We interpolated the data between heel strikes to 100 time points, representing percentage from heel strike to toe-off of the triggering leg. For the resulting interpolated trajectories for the perturbation steps, we subtracted the average of the control steps for the same stance foot. Deviation of the perturbation steps away from the average of the control steps were interpreted as the response to the perceived fall induced by the vestibular stimulus (Δ). For the model corrected foot placement, we fitted a linear regression model relating the foot placement changes for each subject to the changes of lateral position and velocity of the CoM at midstance using the control data (Wang and Srinivasan, 2014). Then for each stimulus step, we used this model to estimate the expected foot placement change based on the CoM state, and subtracted this from the observed foot placement change, resulting in an estimate of the foot placement change due to the visual stimulus (Reimann et al., 2017). We will refer to this model-based estimate as model corrected foot placement change.

### 2.4. Outcome Variables

We hypothesized the lateral ankle mechanism would play a larger role in balance in the *Low* condition, the foot placement mechanism would play a larger role in balance in the *High*. Despite the expected difference in the use of the balance mechanisms, we also hypothesized that the overall shift of the CoM as a result of the response to the balance perturbation would not differ between conditions.

To test our hypothesis about the foot placement mechanism, we analyzed the following five variables that are directly or indirectly related to the first post-stimulus swing leg heel strike: (i) the foot placement is defined as the Δ swing leg heel position relative to the trigger leg heel position. (ii) the model correct foot placement is defined as the measured foot placement value and the foot placement value predicted based on the position and velocity of the CoM at mid-stance using the linear model (see above). (iii) the Δ trigger leg knee internal/external rotation angle. (iv) the Δ swing leg hip internal/external rotation angle. (v) the Δ swing leg hip abduction/adduction angle.

To test our hypothesis about the lateral ankle mechanism, we analyzed the following four variables related to the stance leg lateral ankle activation: (vi) the Δ CoP-CoM distance integrated over the time between the triggering heel strike and the first post-stimulus swing leg heel strike. (vii) the Δ stance leg ankle eversion/inversion angle integrated over the time between the triggering heel strike and the first post-stimulus swing leg heel strike. (viii) the Δ peroneus longus EMG of the stance leg integrated over the time between the triggering heel strike and the first post-stimulus swing leg heel strike. (ix) the Δ tibialis anterior EMG of the stance leg integrated over the time between the triggering heel strike and the first post-stimulus swing leg heel strike.

To test our hypothesis about the push-off mechanism, we analyzed the following two variables related to the stance leg ankle push-off: (x) the Δ plantarflexion angle integrated over the second post-stimulus double stance phase. (xi) the Δ medial gastrocnemius EMG of the stance leg integrated over the first post-stimulus swing phase.

To test the our hypothesis about the overall balance response, we used (xii) the maximum shift of the CoM following the stimulus onset.

### 2.5. Statistical analysis

We confirmed the assumptions of normality and homoscedasticity by visual inspection of the residual plots for the variables related to foot placement, lateral ankle, and push-off mechanisms. Our primary analysis is a group analysis of the kinematic, kinetic, and electromyographical basis for the three balance mechanisms. To test our hypotheses about whether the relative influence of balance mechanisms changes at different cadences, we used R (R Core Team, 2013) and lme4 (Bates et al., 2015) to perform a linear mixed effects analysis. For each outcome variable, we fitted a linear mixed model and performed an ANOVA to analyze the symmetry of the balance response and interaction of the stimulus direction. We used Satterthwaites method (Fai and Cornelius, 1996) implemented in the R-package lmerTest (Kuznetsova, 2017). As fixed effects, we used triggering foot (left/right) to test the symmetry of the response and cadence (*High*/*Low*) to test the difference in response to the balance perturbation in the two conditions. As random effects, we used individual intercepts for subjects. To analyze whether the differences between stimulus and control steps represented by the outcome variables were statistically significant, we calculated the least squares means and estimated the 95% confidence intervals for the intercept of each outcome variable at each level of the significant factor, using a Kenward-Roger approximation (Halekoh and Højsgaard, 2014) implemented in the R-package emmeans (Lenth, 2016). We refrained from approximating p-values for the ANOVA directly in the traditional format, which can currently not be calculated reliably due to the lack of analytical results for linear mixed models (Bates et al., 2015). Results were judged statistically significant in two manners. First, the existence of the use of a variable associated with a balance mechanism in response to the perceived fall is judged statistically significant when the 95% confidence interval did not include zero (bolded in Table 2). Second, the difference in the use of the variables associated with a balance mechanism is judged statistically significant when the 95% confidence intervals for the *Low* and *High* condition did not overlap (highlighted grey in Table 2). We limited the statistical tests to the concrete hypotheses involving the use of the balance mechanisms that we had prior to performing the experiment (Brenner, 2015).

**Table 1:** Results of the ANOVA indicating difference between conditions and which factors have a significant effect on the magnitude of the motor response to the balance perturbation (see Table 2 for statistics on the existence of the mechanism).

| Variable | Fixed Effects | Numerator Df | Denominator Df | F | p |
| --- | --- | --- | --- | --- | --- |
| $\int \Delta$ CoP-CoM | Trigger leg | 1 | 2720 | 0.368 | 0.544 |
|  | Cadence | 1 | 2720 | 275.65 | **<0.0001** |
| $\int \Delta$ Ankle Eversion | Trigger leg | 1 | 2720 | 0.2487 | 0.618 |
|  | Cadence | 1 | 2719 | 542 | **<0.0001** |
| $\int \Delta$ Peroneus Longus EMG | Trigger leg | 1 | 2719 | 3.81 | 0.051 |
|  | Cadence | 1 | 2718 | 10.79 | **0.0010** |
| $\int \Delta$ Tibialis Anterior EMG | Trigger leg | 1 | 2719 | 2.11 | 0.1461 |
|  | Cadence | 1 | 2718 | 314.94 | **<0.0001** |
| $\Delta$ Foot placement | Trigger leg | 1 | 2720 | 1.415 | 0.234 |
|  | Cadence | 1 | 2719 | 267.86 | **<0.0001** |
| Model-Correct Foot placement | Trigger leg | 1 | 2719 | 0.1591 | 0.69 |
|  | Cadence | 1 | 2719 | 1540.43 | **<0.0001** |
| $\Delta$ Knee Rotation | Trigger leg | 1 | 2721 | 0.443 | 0.506 |
|  | Cadence | 1 | 2720 | 3.61 | 0.05715 |
| $\Delta$ Hip Rotation | Trigger leg | 1 | 2720 | 0.1755 | 0.6753 |
|  | Cadence | 1 | 2719 | 325.54 | **<0.0001** |
| $\Delta$ Hip Adduction | Trigger leg | 1 | 2719 | 0.1295 | 0.719 |
|  | Cadence | 1 | 2719 | 125.97 | **<0.0001** |
| $\int \Delta$ Ankle Plantarflexion | Trigger leg | 1 | 2720 | 1.745 | 0.186 |
|  | Cadence | 1 | 2718 | 0.972 | 0.3242 |
| $\int \Delta$ Medial Gastroc EMG | Trigger leg | 1 | 2720 | 5.53 | **0.0188** |
|  | Cadence | 1 | 2719 | 0.903 | 0.342 |
| Max $\Delta$ CoM | Trigger leg | 1 | 2453 | 2.74 | 0.0979 |
|  | Cadence | 1 | 2453 | 7.35 | **0.006** |

**Table 2:**
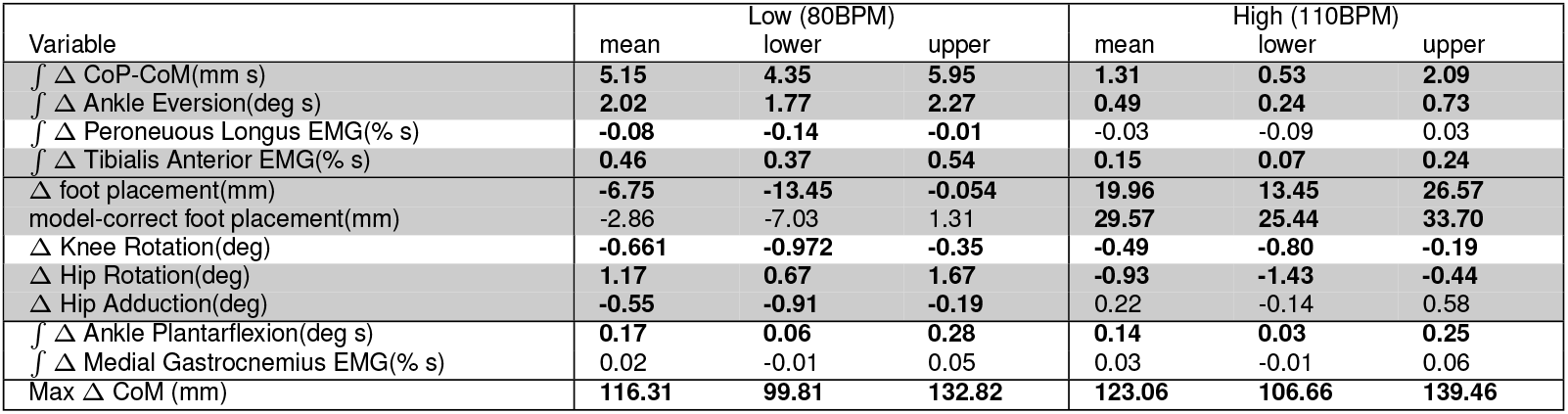
Least-squares means and upper and lower limits of the 95% confidence intervals for each outcome variable for *High* and *Low* cadence conditions using Kenward-Roger approximation. Statistically significant response from zero are marked as bold-face. Statisically significant difference between cadence conditions are marked by grey background.

| Variable | Low (80BPM) |  |  | High (110BPM) |  |  |
| --- | --- | --- | --- | --- | --- | --- |
|  | mean | lower | upper | mean | lower | upper |
| $\int \Delta$ CoP-CoM(mm s) | 5.15 | 4.35 | 5.95 | 1.31 | 0.53 | 2.09 |
| $\int \Delta$ Ankle Eversion(deg s) | 2.02 | 1.77 | 2.27 | 0.49 | 0.24 | 0.73 |
| $\int \Delta$ Peroneus Longus EMG(% s) | -0.08 | -0.14 | -0.01 | -0.03 | -0.09 | 0.03 |
| $\int \Delta$ Tibialis Anterior EMG(% s) | 0.46 | 0.37 | 0.54 | 0.15 | 0.07 | 0.24 |
| $\Delta$ foot placement(mm) | -6.75 | -13.45 | -0.054 | 19.96 | 13.45 | 26.57 |
| model-correct foot placement(mm) | -2.86 | -7.03 | 1.31 | 29.57 | 25.44 | 33.70 |
| $\Delta$ Knee Rotation(deg) | -0.661 | -0.972 | -0.35 | -0.49 | -0.80 | -0.19 |
| $\Delta$ Hip Rotation(deg) | 1.17 | 0.67 | 1.67 | -0.93 | -1.43 | -0.44 |
| $\Delta$ Hip Adduction(deg) | -0.55 | -0.91 | -0.19 | 0.22 | -0.14 | 0.58 |
| $\int \Delta$ Ankle Plantarflexion(deg s) | 0.17 | 0.06 | 0.28 | 0.14 | 0.03 | 0.25 |
| $\int \Delta$ Medial Gastrocnemius EMG(% s) | 0.02 | -0.01 | 0.05 | 0.03 | -0.01 | 0.06 |
| Max $\Delta$ CoM (mm) | 116.31 | 99.81 | 132.82 | 123.06 | 106.66 | 139.46 |

## 3. Results

Subjects adjusted their stepping cadence in order to match the low or high metronome. Figure 2 shows box plots for the step time, step length and velocity for the unperturbed control steps for each cadence condition. Subjects tended to take longer, faster steps as a result of matching a higher frequency metronome (Figure 2A,B). The longer, faster steps resulted in higher overall walking velocities (Figure 2C).

**Figure 2:**
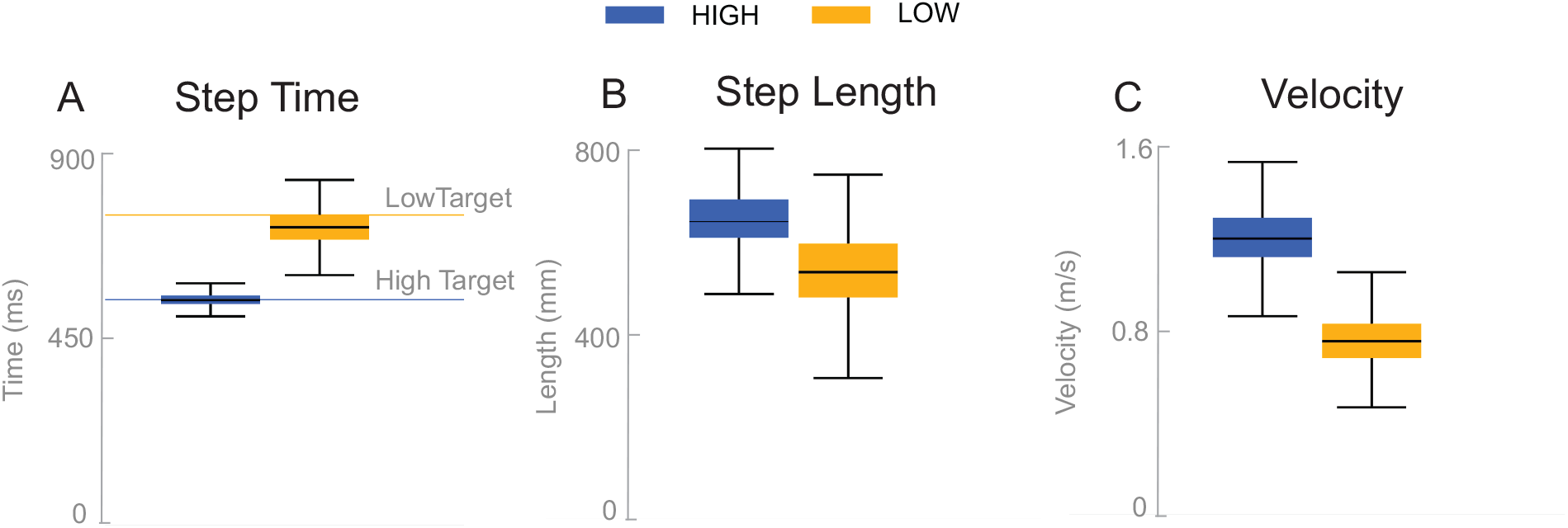
Box plots for the unperturbed control steps. Boxes cover the first to third quartiles, whiskers show the upper and lower adjacents of the data, black lines are the means. Blue boxes correspond to *High* cadence, yellow boxes correspond to *Low* cadence.

Subjects responded to the balance perturbation by shifting their CoM in the direction opposite to the perceived fall direction, i.e. away away from the triggering leg in both conditions (Figure 3). The CoM deviation peaks in the third single stance post-stimulus for the *Low* condition, and peaks in the fourth single stance post-stimulus for the *High* condition. The total CoM excursion was similar in both conditions, with the peak CoM excursion for the *Low* condition 116.31 mm and 123.06 mm for the *High* condition, on average. The peak occurs about the same time in both conditions, ~2 seconds, despite differences in step frequency. In the following paragraphs, we will focus our analysis on the initial responses to the sensory perturbations during the first step and second double-stance period following the triggering heel-strike.

**Figure 3:**
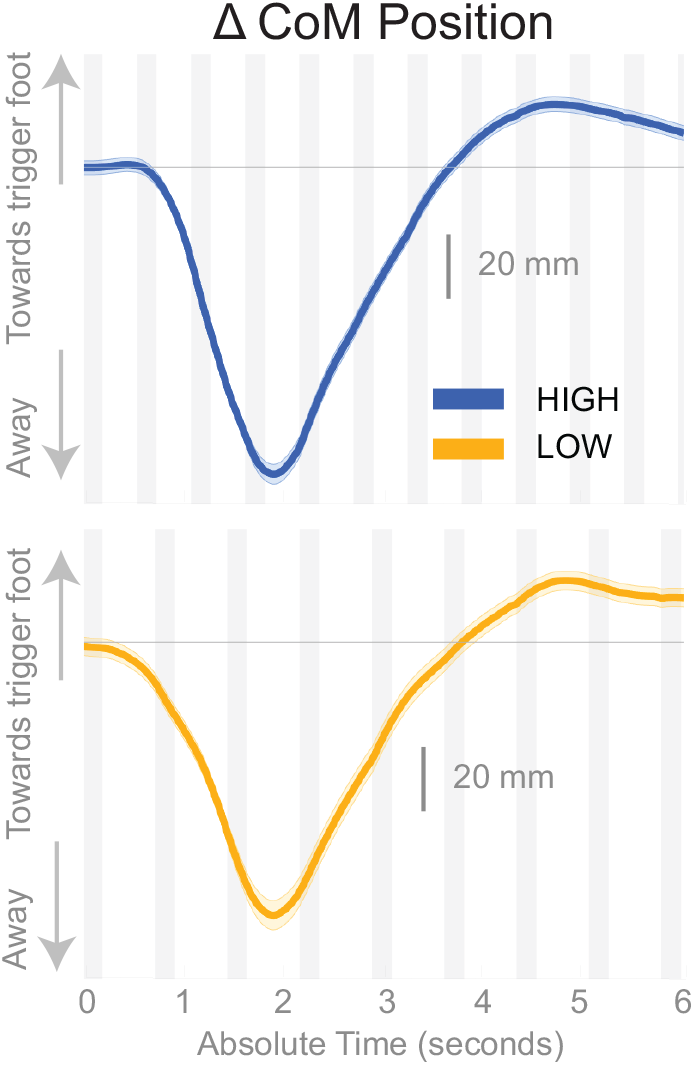
Changes in response to the balance perturbation in the medial-lateral CoM position *High* and *Low* metronome conditions over 12 steps. Curves start at the heel strike triggering the stimulus and end at the twelfth heel strike post-stimulus. The gray, shaded, vertical areas correspond to double stance.

Figure 4 shows the displacement of the CoP relative to the CoM. The CoP shifts in the direction of the perceived fall (see Figure 4), i.e. towards the triggering leg, in both conditions, until the end of single stance. During the second double stance post-stimulus, the CoP moves away from the perceived fall in the *Low* condition, and further towards the perceived fall in the *High* condition.

**Figure 4:**
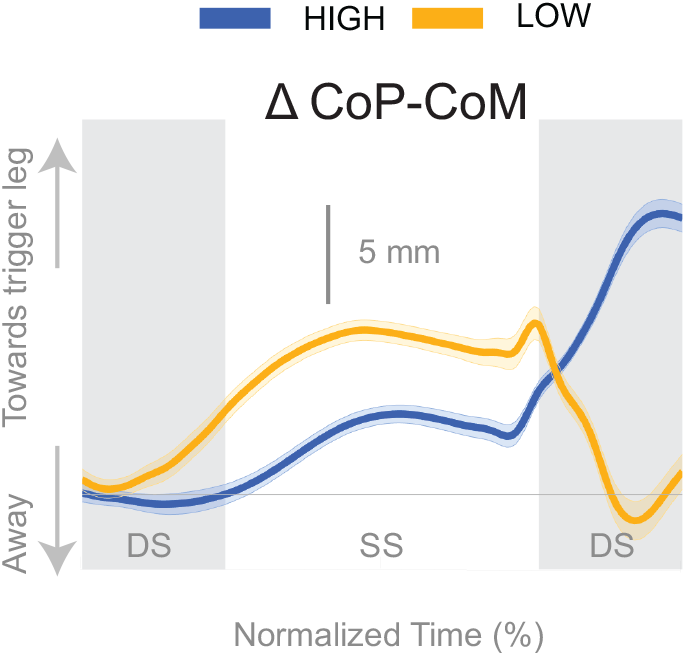
Changes in response to the balance perturbation in the medial-lateral CoP-CoM. Curves start at the heel strike triggering the stimulus and end at the triggering foot toe-off. Blue curves correspond to fall stimuli toward the triggering leg during *High* metronome, yellow curves to fall stimuli toward the triggering leg during *Low* metronome. The gray vertical shaded areas correspond to double stance. DS - double stance, SS - single stance.

The shift of the CoP and CoM are generated by distinct balance mechanisms at different points of the gait cycle. The first mechanism that is able to act is the lateral ankle mechanism. The ankle of the new stance leg inverts during the first post-stimulus step relative to the unperturbed pattern (Figure 5A, Table 2). This ankle inversion change is larger in the *Low* cadence (Figure 5A, Table 2). Activity of the peroneous longus, an ankle everter decreases, beginning in the first double stance for both conditions. Activity of the tibialis anterior, an ankle inverter, increases, beginning in the first double stance post-stimulus for both conditions. These changes are larger in the *Low* condition for the tibialis anterior, but the confidence intervals overlap between the two cadence conditions for the peroneous longus (see Table 2). The timing and magnitude of these changes in kinematics and muscle activation align well with the initial CoP-CoM displacement in the direction of the perceived fall during the first step for both cadence conditions, as shown in Figure 4 (2).

**Figure 5:**
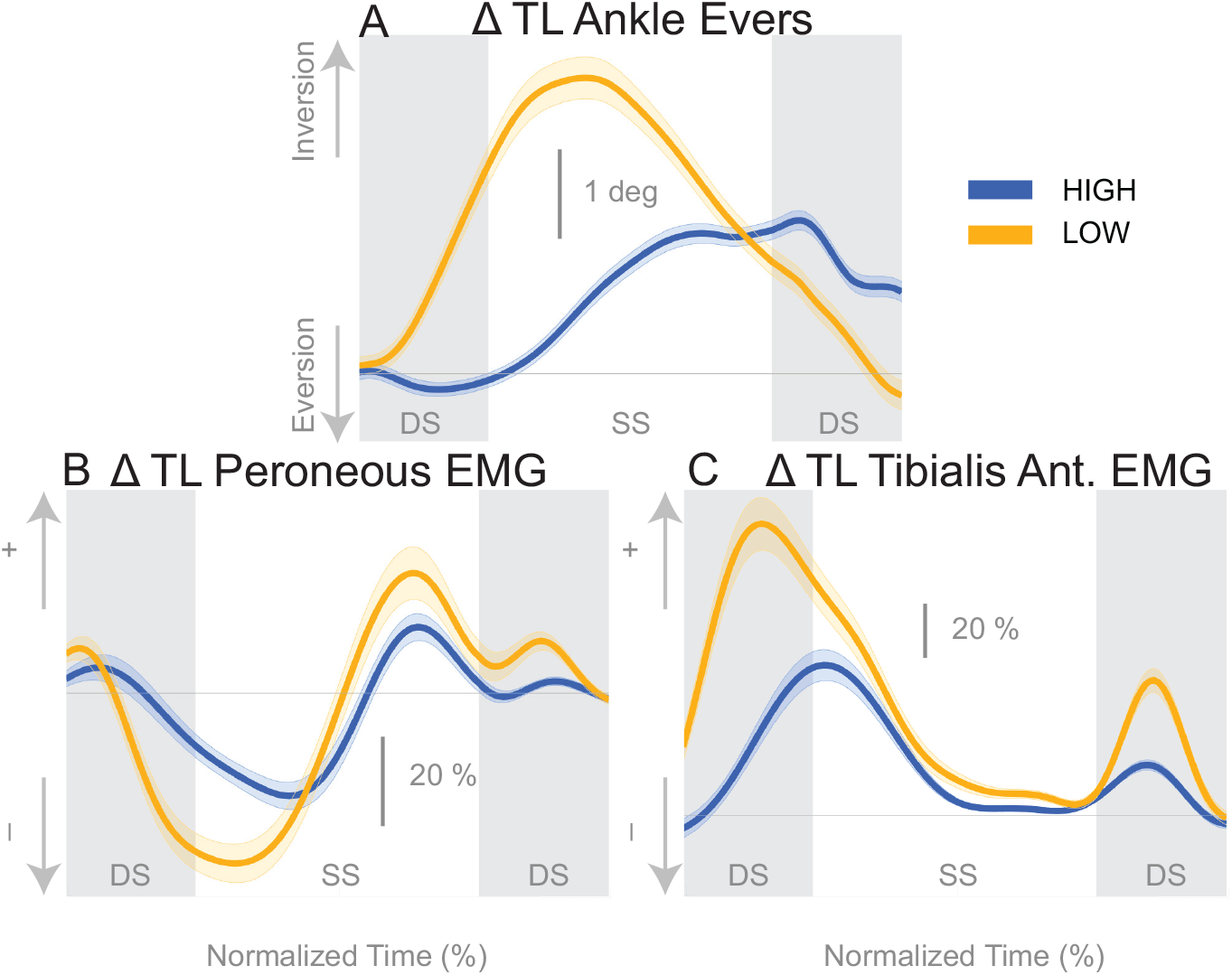
Variables illustrating the use of the lateral ankle mechanism. (A) The joint angle that contributes to the CoP-COM changes; and (B,C) the EMG that contributes to joint angle changes. The baseline represents the average for the control steps. Bold cures indicate the average and the light curves encasing the bold curves indicates the 95% confidence interval. Curves start at the heel strike triggering the stimulus and show the subsequent two steps. Blue curves - *High* metronome, yellow curves - *Low* metronome. DS - double stance, SS - single stance.

At the first post-stimulus heel-strike, subjects shifted their foot placement in the direction of the perceived fall in the *High* condition (Figure 6A), as expected (see Table 2). Contrary to our expectation, subjects tended to shift their foot placement in the opposite direction in the *Low* condition, i.e. away from the perceived fall, although this shift was not statistically significant. Surprisingly, this general pattern was similar for the model-corrected foot placement (Figure 6B), where the shift away from the perceived fall in the *Low* condition is statistically significant, though smaller (see Table 2).

**Figure 6:**
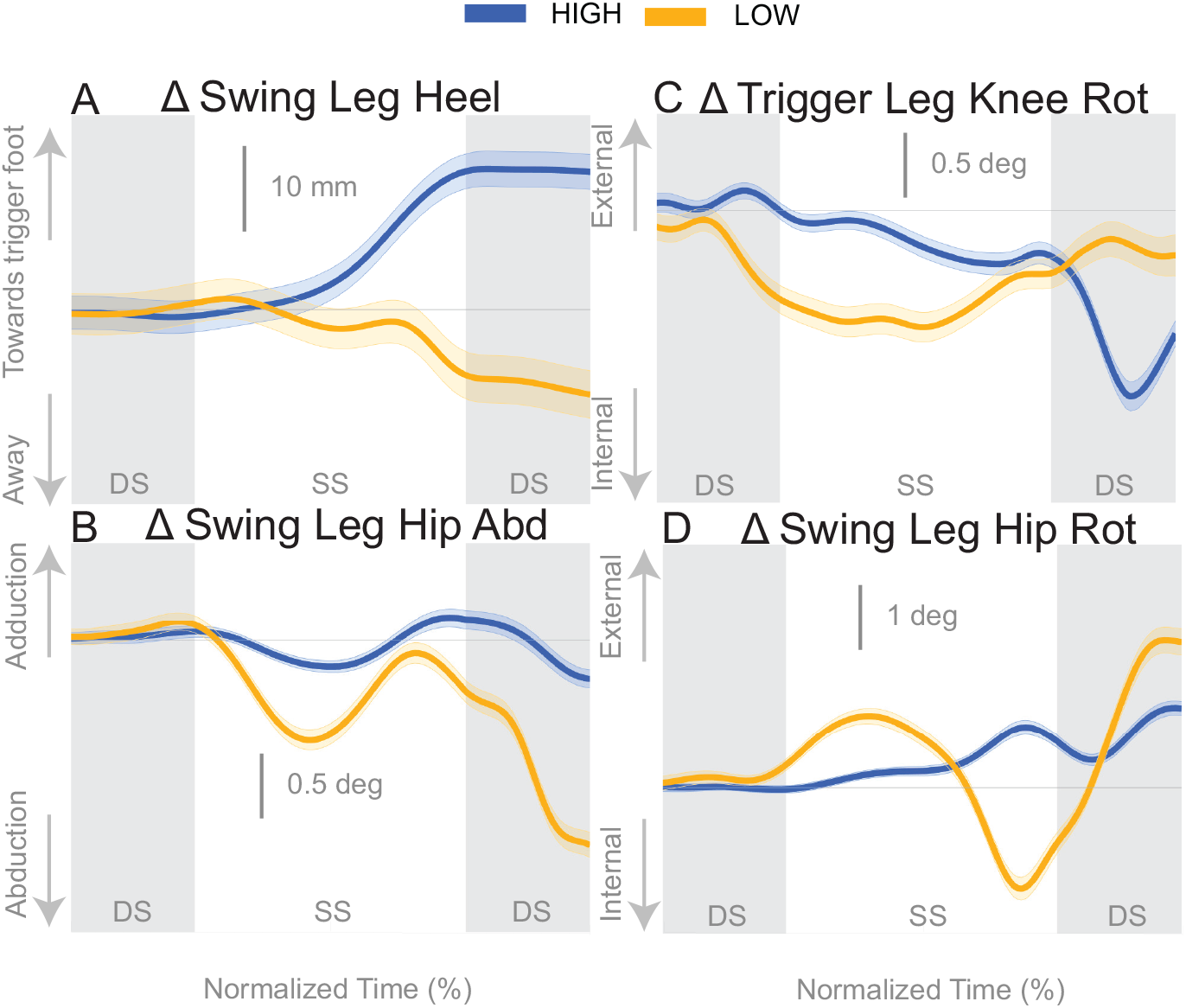
Variables illustrating the use of the change of the foot placement mechanism over time. Changes in response to the balance perturbation in the heel position (A), and joint angles (B,C,D) that contribute to change in heel position. The baseline represents the average for the control steps. Bold cures indicate the average and the light curves encasing the bold curves indicates the 95% confidence interval. Curves start at the heel strike triggering the stimulus and show the subsequent two steps. Blue curves - *High* metronome, yellow curves - *Low* metronome. DS - double stance, SS - single stance.

For the *High* condition, the swing leg hip is slightly adducted upon heel strike (Figure 6B), though not statistically different from control steps (Table 2). A hip adduction modulation would contribute to the swing leg heel moving towards the trigger leg. The *Low* condition produces an abduction of the swing leg hip, allowing for movement of the heel away from the trigger leg. The combination of stance leg knee rotation and swing leg hip rotation yield produce a shift of the swing leg heel in the medial-lateral direction. For example, in the *High* condition, a trigger leg knee internal rotation (Figure 6C) and a swing leg hip external rotation (Figure 6D), in combination, result in a placement of the heel towards the trigger leg. The *Low* condition shows an internally rotated trigger leg knee, but also an internally rotated swing leg hip. The change in leg joint angles support the observed shift in the swing leg heel position.

The push-off mechanism was used for both cadence conditions (Table 2), where an increased trigger leg plantarflexion angle was observed in the second double stance following the triggering heel strike (Figure 8A). However, there was no difference in the plantarflexion modulation between cadence conditions (see Table 2). Similarly, there was no between-cadence difference in the trigger leg medial gastrocnemius EMG (see Table 2).

**Figure 7:**
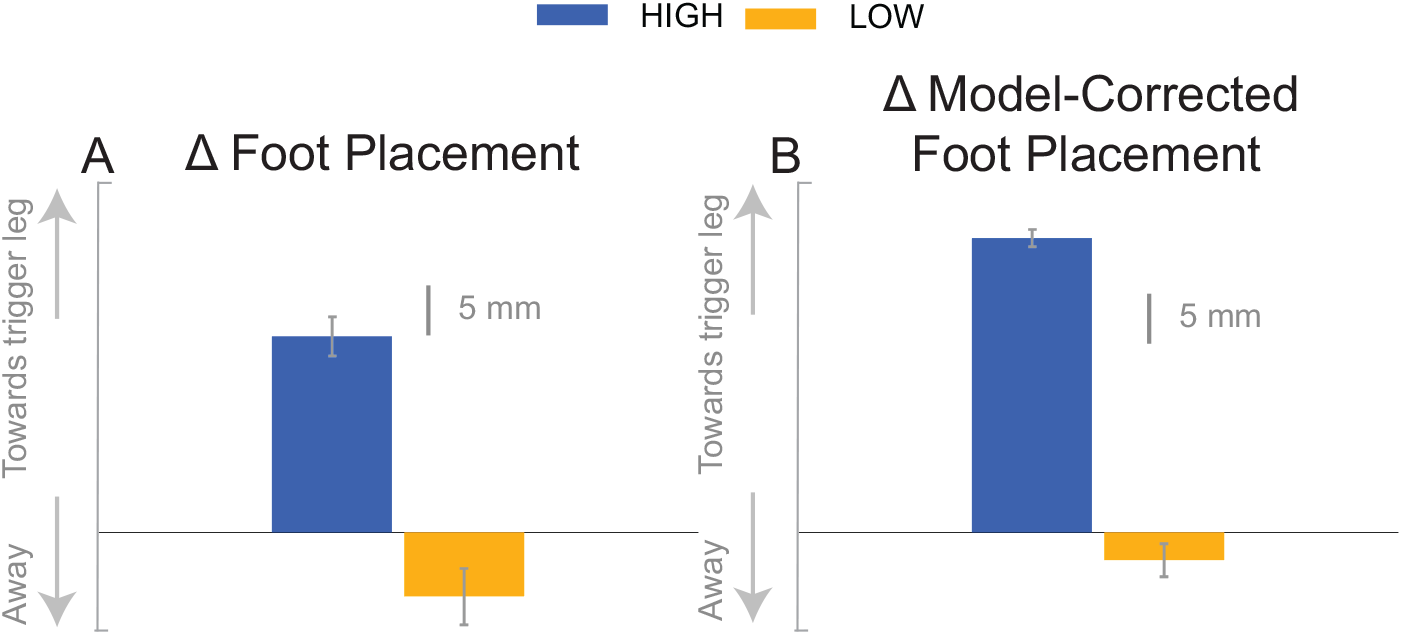
The foot placement response (A) and the model-corrected foot placement (B). The bar graph indicates the mean and the error bar indicates the 95% confidence interval.

**Figure 8:**
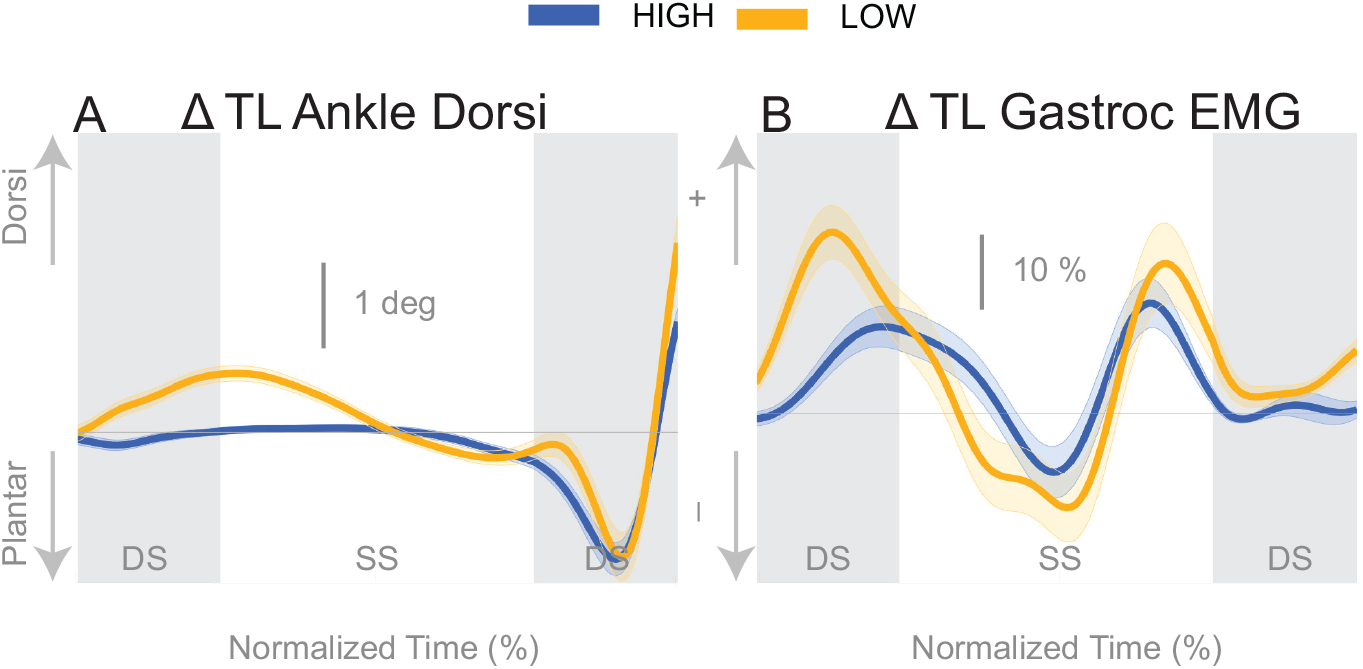
Variables illustrating the use of the push-off mechanism. Changes in response to the balance perturbation in ankle dorsiflexion (A), and the medial gastrocnemius EMG (B). The baseline represents the average for the control steps. Bold cures indicate the average and the light curves encasing the bold curves indicates the 95% confidence interval. Curves start at the heel strike triggering the stimulus and show the subsequent two steps. Blue curves - *High* metronome, yellow curves - *Low* metronome. DS - double stance, SS - single stance.

## 4. Discussion

We studied the changes of the control of balance at different cadences by constraining subjects’ stepping to either *High* or *Low* cadence using a metronome and providing vestibular stimuli that induce the sensation of falling to the side. Subjects responded to the balance perturbations by accelerating their body away from the direction of the perceived fall regardless of the cadence as observed in Figure 3. We limited our analysis to the time interval from the heel-strike triggering the stimulus to the end of the second double stance post-stimulus, which encompasses the three previously identified balance mechanisms of lateral ankle roll, foot placement shift and push-off modulation. We observed changes in kinematics, ground reaction forces, and muscle activation to determine whether the neural control of balance is altered at different cadences. We found distinct changes in the use of balance mechanisms while walking at lower or higher cadences.

We found that the overall effect of the stimulus seems to be invariant of the cadence condition. Neither the time, nor the magnitude of the maximal CoM displacement in the direction of the perceived fall is significantly different between the two cadence conditions. However, the relative use of the mechanisms depends on the cadence. We found that the lateral ankle mechanism played a larger role in the *Low* condition, supported by differences in the CoP-CoM modulation, ankle inversion, and lateral ankle inverter (tibialis anterior) electromyography readings. The ankle inverted to a larger degree in *Low*, indicating that the larger shift of the CoP in the *Low* condition is a result of the use of the lateral ankle mechanism. The increased inversion, along with the prolonged single stance period allows the CoP and CoM to separate for a longer period of time. The combination of decreased peroneous longus muscle activity and increased activity of the tibialis anterior muscle in the *Low* condition compared to the *High* condition, yields an increased inversion angle (Figure 5C). The tibialis anterior is not only an ankle dorsiflexor, but also an ankle inverter, so the combined electromyographical changes to the muscles responsible for ankle inverting suggests the central nervous system is actively generating a larger shift of the CoP under the stance foot in the *Low* condition. The increased use of the lateral ankle mechanism during the *Low* condition makes sense, given the increased duration of single stance in the *Low* condition.

We were surprised by the absence of a foot placement response to the perceived fall in the *Low* condition. We had expected the foot placement shift to be smaller in the *Low* condition, due to an increased and prolonged lateral ankle mechanism response, based on modeling results (Reimann et al., 2017). The lateral ankle mechanism accelerates the CoM, trying to stop the perceived fall, and this CoM shift is sensed by the CNS and invokes a balance response in the opposite direction, which is expected to cancel out the response to the stimulus to some degree. Following Wang and Srinivasan (2014), we fitted a regression model to the relationship between the CoM at midstance and the foot placement shift in the unperturbed reference, controlling for the different cadence conditions (Stimpson et al., 2018). We used this model to estimate the expected foot placement shift based on the normal variability of the CoM movement and calculated the model-corrected foot placement shift by subtracting the model prediction from the observed value. This model-corrected foot placement shift isolates the response to the sensory perturbation, which should be equal between the two cadence conditions. In previous experiments, this model-corrected foot placement shift showed a more consistent response to the sensory perturbation than the uncorrected value (Reimann et al., 2018b). Contrary to this expectation, the model-corrected foot placement change observed at *Low* cadence was significantly lower than at *High* cadence, and even below zero on average, though this was not statistically significant (see Table 2).

The results indicate that all three previously identified balance mechanisms are used to respond to the perceived fall, but the majority of the balance response is shifted to the lateral ankle mechanism when walking with a low cadence. In contrast, when walking at a higher cadence, the foot placement mechanism dominates the balance response. The fact that the push-off mechanism does not differ between conditions indicates that the push-off may not be a critical balance mechanism, but may provide a subtle balance related adjustment at the end of the gait cycle. Unexpectedly, we observed a difference in the dorsiflexion angle between cadence conditions early in the gait cycle, with an increased dorsiflexion in the *Low* condition (see Figure 8A). Counterintuitively, the gastrocnemius EMG is also increased in the *Low* condition early in the gait cycle (see Figure 8B), therefore the dorsiflexion response must be attributed to the increased tibialis anterior activity (see Figure 5C). Such an observation indicates a form of “stiffening” that we plan to systematically analyze in a future publication.

These results support the idea of continuous monitoring of the balance response throughout the gait cycle, and that the balance mechanisms are in fact interdependent (Fettrow et al., 2018). Considering that a change in foot placement has dominated the walking literature as the primary balance response (Bruijn and van Dieën, 2018; Yiou et al., 2017), our results argue against a single balance mechanism for balance control and support the idea of a coordinated response of multiple mechanisms to achieve flexible control of balance while walking. As the vestibular perturbation elicits a perceived acceleration of the CoM in a particular direction, the CNS must shift the CoM an equal amount in the opposite direction to counter the perceived shift. The adjustment can be made continuously throughout the gait cycle, and the total balance response can be viewed as the summation of the balance mechanisms. These results suggest that the lateral ankle and foot placement mechanisms are interdependent while interdependence between the push-off mechanism and the other mechanisms, is weak at best.

These findings will also inform our models of how the balance control system adjusts to altering gait parameters. The foot placement mechanism is the most commonly modeled balance mechanism (Townsend, 1985; Wang and Srinivasan, 2014), but to our knowledge no one has attempted to determine its use at varying cadences. Furthermore, models of human locomotion have difficulty walking at slower speeds (Song and Geyer, 2015). The difficulty with gait speed adjustment may point to limitations in the mechanisms available in these models for the control of balance. The inability for the models to walk slowly could be a result of missing the degree of freedom and control strategy to implement the lateral ankle, foot placement, and push-off mechanism in a coordinated fashion. To allow for a model to maintain walking at different gait speeds, multiple balance mechanisms may be required in order to reproduce as flexible a system as observed with human bipedal gait.

Finally, we speculate that these findings may shed light on the development of preferred walking speeds, particularly in populations that have difficulty with balance. Older adults tend to decrease gait speed with age (Himann et al., 1988; Lauretani et al., 2003), which according to this data set would result in more use of the lateral ankle mechanism. We also know that people with Parkinson’s disease take shorter faster steps (Knutsson, 1972), and although their overall gait speed is diminished, an increase in cadence may shift the majority of the balance response to the foot placement compared to age matched controls. Reasons for why people with Parkinson’s Disease would shift the balance response to foot placement lies in evidence that they have reduced proprioception (Hwang et al., 2016), possibly leading to an inability to sense the CoP under the stance foot. This deficit would make the use of the lateral ankle mechanism unreliable. If a particular balance mechanism is unreliable or cannot be used, the CNS must recruit the available balance mechanisms. In a hypothetical case that the lateral ankle mechanism activation is hindered, it seems logical to increase cadence to rely more on the foot placement mechanism. Thus, we speculate that balance mechanisms play a role in preferred cadence, and possibly gait speed. Future experiments will attempt to directly assess the role of cadence in balance and preferred gait speed in older adults and those with neurological conditions such as Parkinson’s disease.

### 4.1. Limitations

We limited our analysis to the time between triggering leg heel strike and the second double stance post-stimulus because all three balance mechanisms are used within this time frame, but also due to the difficulty interpreting subsequent steps. The response to the perturbation may continue into the next gait cycle (balance perturbation continues for 1000ms), but at some point the CNS responds to the self-inflated fall due to the response to the perceived fall. It is difficult to uncover when precisely the CNS determines the initial response to the balance perturbation is erroneous, but we speculate that the total balance response to the perturbation is similar between cadence conditions due to the similar overall deviation of the CoM in Figure 3.

Our current results show that higher cadence results in faster gait speed, at least for healthy young adults walking on a self-paced treadmill. The relationship between gait speed and cadence is non-trivial, making it problematic to unpack the effects of cadence versus gait speed. The convergence to a faster gait speed when adopting a higher cadence may be related to metabolic efficiency, but this is an avenue far removed from the focus of the current work. To our knowledge this relationship has not been explored.

Our current experimental methods may lead to limitations in the presentation of the push-off mechanism. We refer to the change in ankle plantarflexion in response to the balance perturbation as push-off, because to date this is the best term to describe what we believe the change in ankle plantarflexion does. Due to the experimental design, we do not have reliable ground reaction forces and moments. Providing balance perturbations, in addition to instructions to “walk normally”, leads to many situations where one foot is on two force plates, or during double stance, two feet are on one force plate. These situations make it difficult to determine the ground reaction forces and moments associated with the change in ankle plantarflexion angle.

Another technical consideration in the current methodology is the heel strike identification method. Different walking speeds in the cadence conditions produced different vertical heel trajectory profiles which required intervention by the experimenter (see Methods). Increasing the vertical threshold created a longer period of time from threshold crossing to actual heel strike for the *Low* condition. Therefore, the stimulus for the *Low* condition was triggered on average ~175ms prior to heel strike, and the stimulus for the *High* condition was triggered ~110ms prior to heel strike, coincidentally corresponding to ~20% of the time between heel strikes for each condition. The earlier trigger in the *Low* condition may partially explain the earlier response time in the CoP-CoM (Figure 4) and corresponding EMG for the lateral ankle mechanism (Figure 5B,C), but not the amplitude.

## 5. Conclusion

We investigated whether individuals altered the control of balance if they were asked to step at different frequencies while walking on a self-paced treadmill as they periodically received a vestibularly-induced sensation of a fall to the side. The current findings support the idea that balance mechanisms are coordinated to produce an overall balance response. In the *Low* condition, the lateral ankle mechanism plays the primary role in the overall balance response, while the foot placement mechanism is not observed. When cadence is decreased (*Low*), single stance is longer, providing more opportunity to modulate the CoP through ankle roll, and diminishing the need for a change in foot placement to maintain upright posture. In the *High* condition, with shorter single stance duration, the lateral ankle mechanism is less effective and the balance response shifts to reliance on the foot placement mechanism. These findings suggest lateral stability is not dependent on cadence or gait speed, but the method of obtaining stability is altered with cadence, providing evidence of a flexible neural control scheme that adapts to changing constraints while walking. Moreover, such findings provide insight into adoption of preferred gait parameters, particularly in populations who have drastically altered gait speed or cadence such as older adults or people with Parkinson’s disease.

